# Mixed data analysis detected Endocrine Fibroblast Growth Factors (FGF19, FGF21, and FGF23) as Prognostic and Diagnostic Markers of Colorectal Neoplasia and Carcinoma

**DOI:** 10.1101/2023.06.27.546716

**Authors:** Leili Rejali, Moein Piroozkhah, Mana Jahanbin, Pooya Jalali, Binazir Khanabadi, Elahe Daskar Abkenar, Mehdi Tavallaei, Mahsa Saeedi Niasar, Mehrdad Hashemi, Amir Sadeghi, Zahra Salehi, Ehsan Nazemalhosseini-Mojarad

## Abstract

**Background:** Endocrine fibroblast growth factors (e*FGFs*) play important roles in various cellular signaling processes such as development and differentiation. These genes were also found to be significantly related to several cancer. However, little is known about the role of e*FGFs* in colon neoplasia and colon adenocarcinoma (COAD).

**Methods:** We performed systematically and comprehensively investigated the gene expression, DNA methylation, prognostic significance, genetic alteration, co-expressed genes, protein-protein interaction, small molecules pathway, and drug interactions of *eFGFs* based on the TIMER2.0, GEPIA2, UALCAN, OncoDB, cBioPortal, LinkedOmics, STRING, SMPDB, htfTarget, mirTarBase, circBank and DGIdb databases. Ultimately, the correlations of *eFGFs* expressions between polyp and COAD tissues compared to normal mucosa were validated using qRT-PCR as a cross-sectional part of our study.

**Results:** The results indicated that *eFGFs* are highly expressed in COAD, and abnormal gene expressions may be related to promoter methylation. In this matter, methylation analysis revealed promotor hypermethylation of *FGF19* and *FGF21*. Conversely, *FGF23* was shown to have a tendency for promotor hypomethylation. Moreover, hypermethylation of *FGF21* and *FGF23* and downregulation of *FGF23* were found to be detrimental to the survival of COAD patients. KEGG pathway analyses indicated that the co-expressed genes of *eFGF* family members were mainly related to the regulation of the actin cytoskeleton and, more notably, in Ras signaling, PI3k-Akt signaling, Rap1 signaling, and cancer pathways. Based on qRT-PCR results, *FGF21* was significantly overexpressed in the colon polyps compared to normal mucosa. Additionally, RNA expression of *FGF21* and *FGF23* was markedly elevated in adenomatous polyps as opposed to hyperplastic polyps.

**Conclusion:** Collectively, these findings reveal the critical roles of *eFGFs* in COAD tumorigenesis and suggest *eFGF* family members as promising prognostic and diagnostic markers for CRC as well as discriminating markers for high-risk from low-risk polyps.

## 1. Introduction

In the United States, colorectal cancer (CRC) is projected to be the second most significant cause of cancer-related deaths in 2023 (1). Nonetheless, new studies suggest a decline in CRC death rates during the previous few years. This appears to be mainly attributable to the implementation of effective screening programs that have resulted in increased detection of premalignant polyps (2). Polyps could be divided into three histological categories: adenomatous (60–70%), serrated (including hyperplastic) (10–30%), and other (10–20%). It’s commonly recognized that CRCs are usually caused by adenomatous polyps (3). Hyperplastic polyps (HPP) are benign, small in size, and noncancerous lesions (4). For the time being, the pathologic diagnosis of HPP and AD depends mainly on histological investigation to detect epithelial morphological alterations. Yet, due to the absence of precise diagnostic biomarkers, it is not always possible to distinguish between them efficiently (5).

Fibroblast growth factors (*FGFs*) are members of the peptide growth factor family that share several roles, such as cell differentiation and growth, embryonic development, angiogenesis, wound healing, and metabolic control (6). Based on their biological activities, evolutionary relationships, and sequence homology, *FGFs* are categorized as intracrine, paracrine, or endocrine. *FGF19*, *FGF21*, and *FGF23* are endocrine factors with modest mitogenic activity. These genes influence physiological processes through interactions with fibroblast growth factor receptors (*FGFR*), but across a greater distance as hormones (7, 8). Endocrine *FGFs* (*eFGF*s) are primarily related to controlling cellular metabolic functions, notably the homeostasis of energy, bile acid, lipids, glucose, and minerals (phosphate/active vitamin D) (9, 10).

*FGF19*, one of the hormone-like *FGFs*, is commonly amplified and overexpressed in several types of malignancies (11–13). Increased levels of *FGF19* cause pre-neoplastic changes in the colon by inhibiting bile acid production and disrupting cholesterol metabolism in the liver (14). The *FGF21*’s pleiotropic metabolic effects in response to various physiological and pathological stressors have garnered considerable attention over the past two decades (15). All in all, upregulation of *FGF21* has been reported in several kinds and stages of cancer, including non small cell lung cancer (16), endometrial cancer (17), and CRC (18), highlighting the biomarker significance of *FGF21* in cancer diagnosis and therapy (15). *FGF23* is a bone-derived hormone like *FGF* that primarily regulates vitamin D and phosphorus metabolism (19). According to mounting evidence, *FGF23* is thought to be essential for some processes that contribute to the development and spread of cancer (20). *FGF23* is especially important for kinds of cancer that predominantly impact the bone (like multiple myeloma) or are characterized by bone metastases (21). New research indicates that malignancies such as CRC (22) and endometrial cancer (17) can also express *FGF23* in an executive manner.

Since *eFGFs* have been shown to have a novel and complicated interaction with metabolism and mitogenic activity, it stands to reason that they may also be closely tied to the development of CRC, as was previously described. Therefore, we explored the function of *eFGF* in colorectal neoplasia and carcinoma using mixed data analysis, including RNA sequencing (RNA-Seq) data derived from multiple high-throughput online databases followed by experimental validation based on qRT-PCR. Taken together, these results provide light on the relationship between the diagnosis and prognosis of colorectal neoplasia and carcinoma and *eFGF* family members.

## 2. Materials and Methods

To better understand the significance of endocrine *FGFs* in COAD, first, by the use of high throughput databases, we measured these gene expressions in pan-cancers. Next, we analyzed the mRNA expression and methylation of *eFGF* in COAD patients and assessed their associations with different clinicopathological characteristics as well as overall survival rates. Examination of genetic alteration of *eFGFs* Across Different COAD Studies was the other part of our study. Thereafter, we identified the co-expressed genes of *eFGF* in COAD and their functional enrichment analysis. We also constructed a protein-protein interaction network of *eFGF* genes. We have successfully constructed gene regulatory networks that consist of gene-TFs and competing endogenous RNA (ceRNA) networks. The first portion of the study was completed by examining the small molecular pathways of the genes mentioned above. Eventually, the bioinformatics analysis results were validated by qRT-PCR. Figure 1 shows a schematic illustration of the analysis and experiments we carried out to account for the role of *endocrine FGF* genes in colorectal carcinogenesis.

**FIGURE 1.**
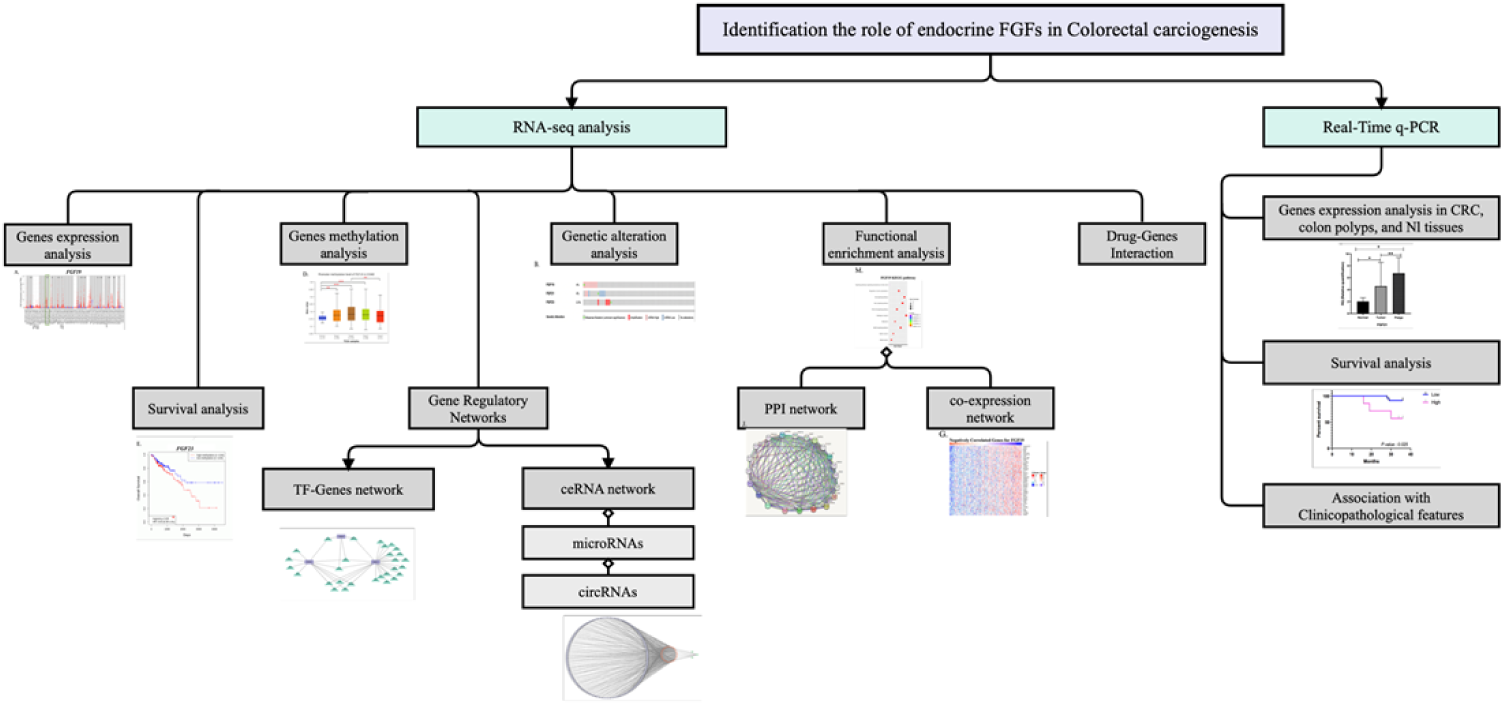
A schematic illustration of the bioinformatics analysis and experimental studies in our investigation.

### 2.1. High-Throughput analysis of endocrine FGFs pattern in CRC

#### 2.1.1. Transcriptional Expression and Clinicopathological Correlations

Initially, we gathered *eFGFs* expression patterns in 33 different cancer tissues compared to the normal adjacent tissue (NAT) and normal healthy donors using the TIMER2.0 (https://cistrome.shinyapps.io/timer/) and GEPIA2 (http://gepia2.cancer-pku.cn/#index) databases, respectively (23, 24). Further, the expressions of *eFGFs* were compared in COAD and para-cancerous tissue samples using the UALCAN (https://ualcan.path.uab.edu) database (25). This database also uncovered associations between clinicopathological features and *eFGFs*. We subsequently established a correlation between the expression of *eFGFs* and the tumor-node-metastasis (TNM) stage of the tumor via GEPIA2.

#### 2.1.2. Analysis of the Promoter Methylation of Endocrine FGF Genes in COAD Patients

In addition, we used UALCAN to compare the methylation levels of *FGF19*, *FGF21*, and *FGF23* promoters across COAD tissues to NAT and other clinicopathological factors. The beta value in our study ranges from 0 (not methylated) to 1 (fully methylated), representing the degree of DNA methylation. Diverse cut-off beta values were applied to distinguish between hypermethylation (beta value: 0.7-0.5) and hypomethylation (beta value: 0.3-0.25) (26, 27); and differences were considered to be significant at a *P-value* of less than 0.05.

#### 2.1.3. Assessing the Association of the Endocrine FGF Family Members’ Expression and Methylation with COAD Patients’ Overall Survival

The OncoDB (http://oncodb.org/) server was used to investigate the relationship between *eFGF* expression or methylation in COAD patients and the overall survival (OS) (28). The results were examined using the *P*-value and HR of COAD patients regarding the differential level of *eFGFs* expressions and methylations depicted in the Kaplan-Meier (KM) plot of survival analysis.

#### 2.1.4. Examination of Genetic Alterations in Endocrine FGF Genes across COAD Studies

Cancer genomic data for multidimensional in silico analysis are easily accessible through cBioPortal (https://www.cbioportal.org/) (29). We analyzed the genetic mutations of *eFGFs* in COAD using this database. The OncoPrint summary of the overall mutations of *eFGFs* in COAD studies was reviewed. OncoPrint of cBioPortal displays mutation types and their corresponding proportions of queried genes in selected studies. Lollipop figures were also generated for *eFGF* genes in colorectal adenocarcinoma (TCGA, Firehose Legacy) studies to align mutations with their protein domains. Throughout the analysis on the cBioPortal server, the parameter values were left at their default settings.

#### 2.1.5. Identification of the Co-Expressed Genes, PPI Construction, and KEGG pathway analysis of Endocrine FGFs in COAD Patients

LinkedOmics (http://www.linkedomics.org), which provided the data from the TCGA-COAD cohort, was used to identify the co-expressed genes of *FGF19*, *FGF21*, and *FGF23* (30). This study visualized Co-expressed genes by volcano plot and heat map (false discovery rate, |FDR|<0.01). A protein-protein interaction network (PPI) of these co-expressed genes was also built using the STRING (https://string-db.org) database (31). Then, via the Enrichr database (https://maayanlab.cloud/Enrichr/) (32), we carried out KEGG pathway analysis of the co-expressed *eFGFs* genes in COAD samples. The “ggplot2„ R package was used for data visualization.

#### 2.1.6. Small Molecules Pathway

This study investigated the small molecule pathways related to *eFGFs* using the SMPDB (https://smpdb.ca) database. The human metabolism, pharmacological action, drug metabolism, physiological activity, and metabolic disease pathways may all be seen in great detail with the help of the SMPDB database (33).

#### 2.1.7. Transcription factors

To detect the Transcription factors (TFs) regulating *eFGF* family members, we employed hTFtarget database (http://bioinfo.life.hust.edu.cn/hTFtarget#). Only those TFs that target *eFGFs* in colon tissues were selected for further analysis. The TF-Genes regulatory network was built and mapped utilizing the Cytoscape tool (version 3.9.1).

#### 2.1.8. The circRNAs-microRNAs-mRNAs network

We used the miRTarBase database (https://mirtarbase.cuhk.edu.cn) to find the miRNAs linked to eFGF genes. Then, we accessed the circBank database (http://www.circbank.cn) to discover the circRNAs that target obtained miRNAs. We sorted the lists of circRNAs-related miRNAs based on ’Total Value’ and chose thirty circRNAs with the highest ’Total value’ for each miRNA for further examination. Subsequently, we constructed ceRNA networks between mRNAs, miRNAs, and circRNAs using Cytoscape tool (version 3.9.1).

#### 2.1.9. Drug-Gene Interaction

The drug-gene interaction database (DGIdb) database (https://dgidb.org/) was utilized to discover the associated medicines for the *eFGF* family members. This database presents information on medications that target hub genes from a number of trustworthy databases and medical literature (34).

### ***2.2.*** Real-time PCR analysis of endocrine FGFs pattern in colorectal cancer and Neoplasia

#### 2.2.1. Patients’ Tissue Specimens and Clinical Data Collection

Finally, a total of 54 colon polyp tissue samples, including 26 AD and 28 HPP, with a statistical power of 90% and a 5% error rate. The mean ± SD of endocrine FGF gene expressions in adenomatous polyps was 0.85 ± 0.56, while in hyperplastic polyps, it was 0.37 ± 0.56, were taken from 100 patients who had a colonoscopy or flexible sigmoidoscopy screening and polypectomy at Taleghani Hospital, Shahid Beheshti University of Medical Sciences (Tehran, Iran) from Jan 2021 to Jan 2022.

In addition, 30 CRC tissues were collected together with clinical data from patients diagnosed with primary CRC and who had undergone surgery at the abovementioned facility. Thirty healthy, normal colon mucosa samples were collected from volunteer individuals after routine colonoscopies. No patients had previously had treatment for cancer with chemotherapy, radiation, targeted therapy, or any other active severe disease. Those who reported a personal or family history of previous cancer or colon polyps were also excluded. The authors had access to identifying information during data collection, but all identifying information was kept confidential and stored separately from the study data. Identifying information was removed and replaced with unique study numbers before data analysis. The authors did not have access to any identifying information after data collection. To determine the postoperative CRC pathological stage, we utilized the TNM grading method developed by the American Joint Committee on Cancer (35). Polyp-type classifications were determined by an expert pathologist based on the polyp’s histology. Written informed consent was obtained from all patients, and the research was approved by the Ethics Committee of Taleghani Hospital at Shahid Beheshti University of Medical Sciences (protocol number: IR.SBMU.RIGLD.REC.1396.180).

#### 2.2.2. RNA Isolation, cDNA Synthesis, and Real-Time Polymerase Chain Reaction (qRT-PCR)

We used TRIzol reagent to extract total RNAs from 116 tissue samples (56 polyps, 30 CRC, and 30 normal colon mucosa) (YTzol pure RNA Yekta Tajhiz Iran). The total RNAs were then reverse transcribed into complementary DNA (cDNA) using cDNA Reverse Transcription kits (Yekta Tajhiz kit Cat: YT4500, Iran) in accordance with the manufacturer’s instructions. Eventually, qRT-PCR was carried out utilizing the SYBR Green-based RT-PCR Master Mix with the cDNA as the template (Yekta Tajhiz, Iran). Relative gene expression was calculated by the 2^−ΔΔ*C*^ cycle threshold method. Detailed information for the primers in Supplementary Table S5 can be obtained.

#### 2.2.3. Statistical analysis

With GraphPad Prism 5.0, data were plotted, and statistical analyses were conducted (GraphPad Software, Inc). The REST program determined fold changes in the gene expression (36). The chi square test was used to analyze the correlations between gene expression and clinicopathological characteristics. Student’s t-test was used to compare polyp or CRC tissues to normal tissues, and one-way analysis of variance (ANOVA) was used to compare different groups. Survival analysis was performed using the log-rank test. Missing values were not inferred. Statistical significance was assumed for *P-values* and *log-rank p* of 0.05 or below.

## 3. Results

### 3.1. The Expression Analysis of eFGFs in CRC Patients

To explore the transcriptional expression of *eFGF* family members in COAD, we analyzed 33 types of cancer tissues and the NAT using the TIMER2.0 database. As shown in Figures 2A, 2B, and 2C, the mRNA expressions of *FGF19*, *FGF21*, and *FGF23* were significantly upregulated in COAD tissues compared to NAT. Besides, the *eFGF*s were also considerably overexpressed in rectum adenocarcinoma, stomach adenocarcinoma, and lung squamous cell carcinoma (Fig. 2A-C). However, when we analyzed the GTEx portal of pan-cancer via the GEPIA2 database, no significant differences were found in the expression of the aforementioned genes between COAD tissues and healthy individuals (Fig. 2D-F). Moreover, comparing the expression of *eFGF* family members based on the TCGA data by UALCAN declared obviously higher expression of *FGF19* and *FGF23* in COAD tissues than NAT (*P-value*= 2.59E-04 and 3.76E-04; respectively). Nevertheless, no statistically significant difference was found for *FGF21* (*P-value* > 0.05) (Fig. 2G-I).

**FIGURE 2.**
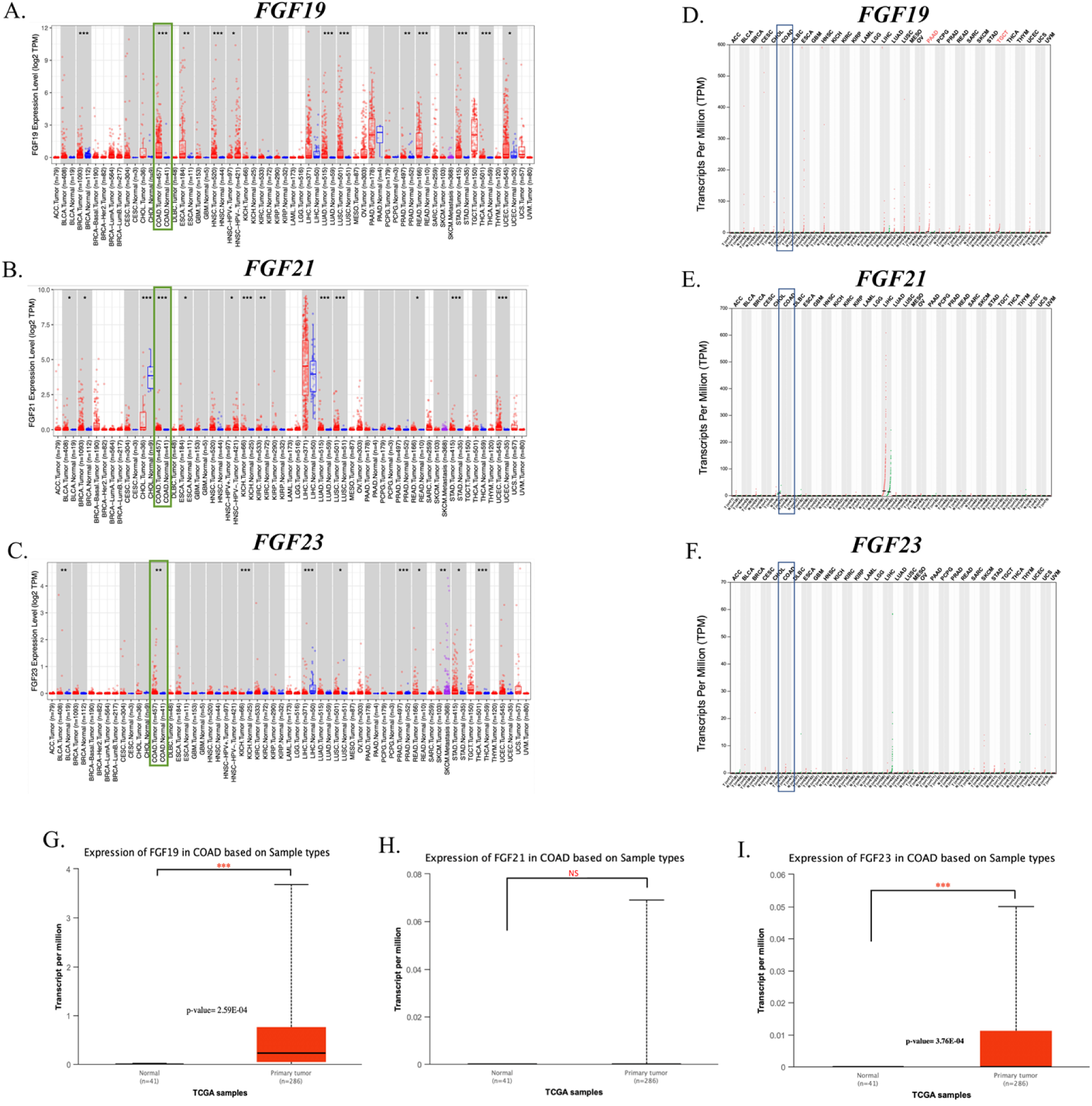
Transcriptional expression of endocrine *FGFs* family member analysis. The mRNA levels of *eFGF* family members in 33 types of cancers from the TIMER2,0 database **(A-C)** and GEPIA2 database (**D-F**). **(G-I)** The expression of *eFGF* family members between COAD and normal tissues from the TCGA. (NS: *P*-value > 0.05, *: *P*-value < 0.05, **: *P*-value < 0.01, ***: *P*-value < 0.001). NS; Non-significant

### 3.2. Association of Transcriptional Expression of eFGFs and the Clinical Parameters in Colon CRC Patients

The results of UALCAN’s inquiry on the correlation between clinicopathological characteristics and mRNA expression of the *eFGF* family revealed that *FGF19* and *FGF23* expressions were substantially and positively correlated with patients’ sex, age, cancer stages, nodal metastasis, and tumor histology (Fig. 3A-J). *FGF19* expression was elevated in stages I, II, and IV compared to normal tissues (*P-value* = 1.61 E-02, 1.3 E-02, and 7.55 E-04, respectively) (Fig. 3E); likewise, *FGF23* expression level was substantially correlated with stages II, III, and IV (*P-value* = 2.43 E-02, 4.62 E-02, and 4.71 E-02, respectively) (Fig. 3F). Based on the status of nodal metastases, from no regional nodal spread (N0) to several degrees of nodal spread (N1-N2), *FGF19* and *FGF23* expressions were significantly higher in N0 (*P-value* = 3.1E-03 and 1.10E-02) and N1 (*P-value* = 8.06E-03 and 4.84E-02) compared to normal colon tissue (Fig. 3G and H). We also investigated the relationship between the histology of COAD tumors and the mRNA expressions of *FGF19* and *FGF23*. The mRNA expression of *FGF19* was significantly higher in adenocarcinoma than in normal tissues (*P-value* = 6.62 104) (Fig. 3I). Additionally, adenocarcinoma (*P-value* = 2.32 E-03) and mucinous adenocarcinoma (*P-value* = 3.12 E-02) overexpressed *FGF23* compared to normal colon tissue (Fig. 3J). Despite this, the UALCAN analysis found no correlation between the expression of *FGF21* and any of the clinicopathological parameters of CRC patients (Fig. S1). Moreover, GEPIA2 demonstrated that among the members of the *eFGFs*, only *FGF19* expression is connected to the TNM stages of COAD (Fig. 3K-M).

**FIGURE 3.**
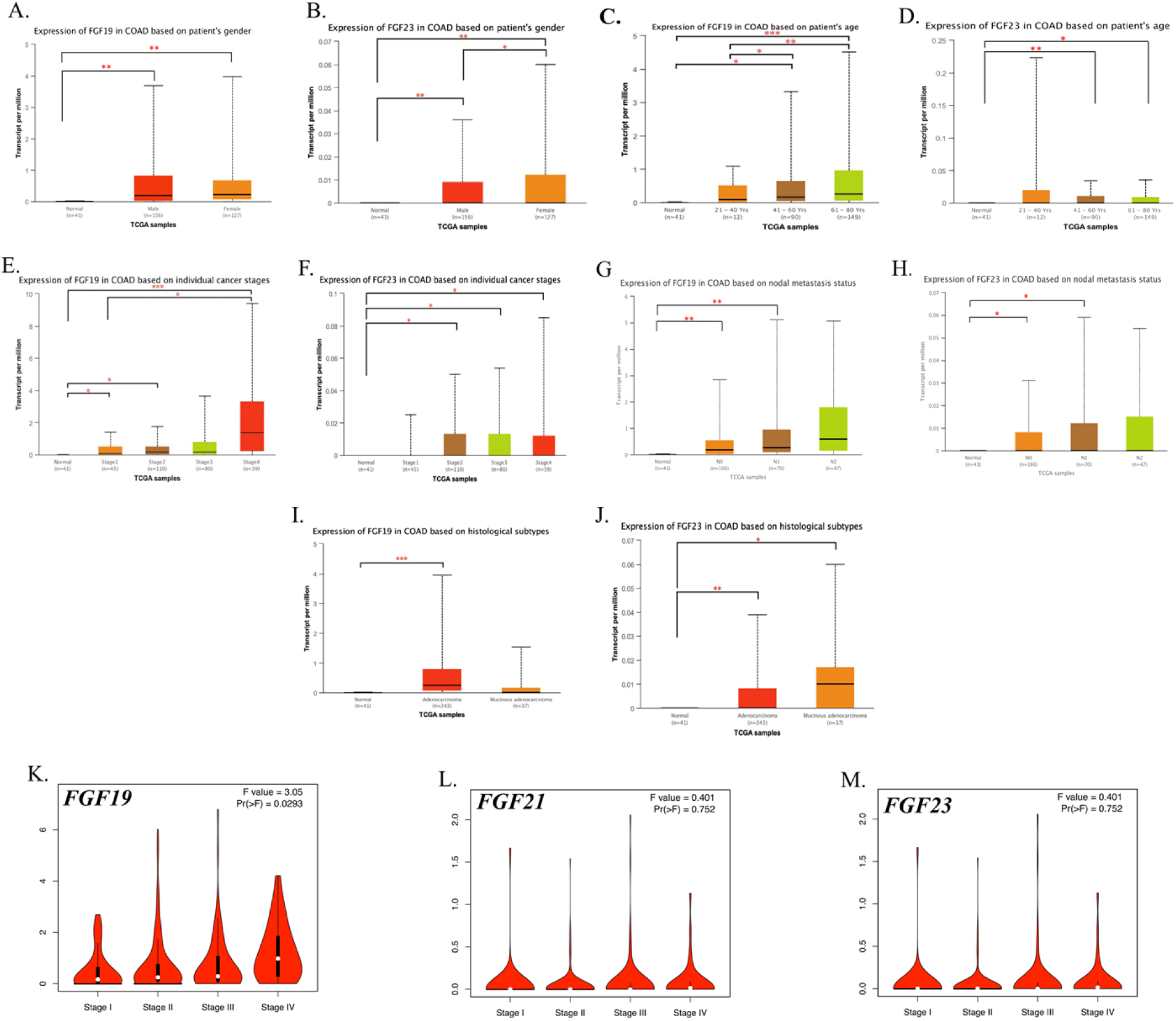
Association between transcriptional expressions of *FGF19 and FGF23* and clinical parameters. The transcriptional expressions of *FGF19 and FGF23* associated with patients’ gender **(A and B)**, age **(C and D)**, cancer stages **(E and F)**, nodal metastasis status **(G and H)**, and tumor histology **(I and J)** based on the UALCAN database. **(K-M)** Association between pathological tumor stages with *FGF19* **(K)**, *FGF21* **(L)**, and *FGF23* **(M)** based on the GEPIA2 database. (NS: *P*-value > 0.05, *: *P*-value < 0.05, **: *P*-value < 0.01, ***: *P*-value < 0.001). NS; Non-significant.

### 3.3. Analysis of eFGFs Promotor Methylation levels in the CRC Patients

It is generally recognized that DNA methylation, a critical form of epigenetic regulation, modifies the state of gene expression (37). In this regard, using the UALCAN database, we further analyze the *eFGFs* promoter methylation levels of COAD. We discovered that in COAD, *FGF19*, and *FGF21* promoters were hypermethylated (*P-values* of 3.43 E-11 and 3.44 E-15; respectively) (Fig. 4A and B). Nonetheless, there was a tendency toward hypomethylation of the *FGF23* promoter (*P-value* = 1.39 E-08) (Fig 4C). Above all, promoter methylation of *eFGFs* was associated with several clinicopathological characteristics, such as tumor TNM stage (Fig. 4D–F) and tumor histology (Fig. 4G–I).

**FIGURE 4.**
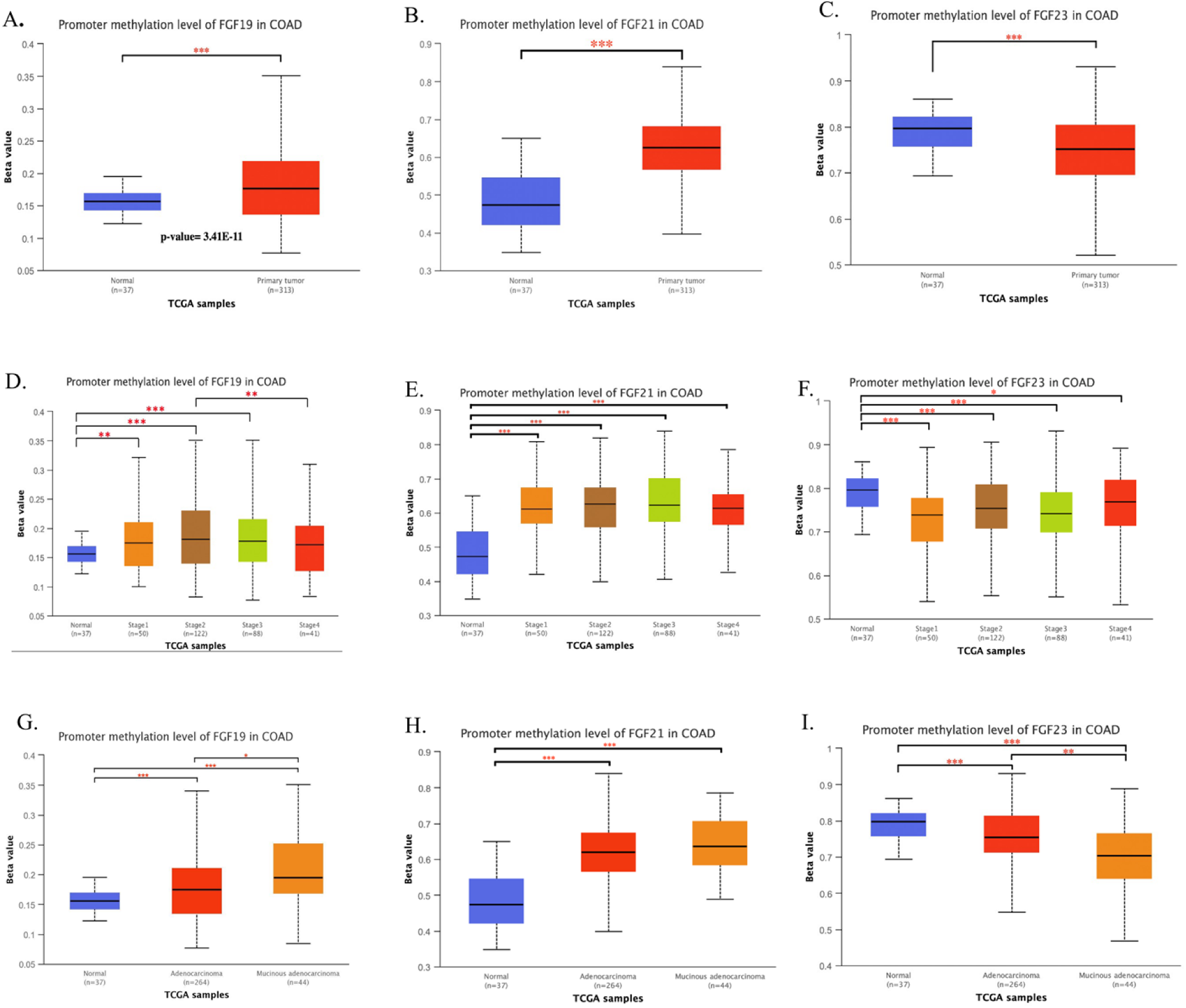
Promoter methylation levels of endocrine *FGFs* in COAD through UALCAN. The variables are sample types **(A-C)**, cancer stages **(D-F)**, and tumor histology **(G-I)**. The Beta value indicates the level of DNA methylation ranging from 0 (unmethylated) to 1 (fully methylated). Different beta value cut-off has been considered to indicate hyper-methylation [Beta value: 0.7 - 0.5] or hypo-methylation [Beta value: 0.3 - 0.25]. (NS: *P*-value > 0.05, *: *P*-value < 0.05, **: *P*-value < 0.01, ***: *P*-value < 0.001). NS; Non-significant.

### 3.4. Survival Analysis of eFGF Family Members Expression and Methylation in the CRC Patients

Furthermore, utilizing the OncoDB database, a Kaplan-Meier plotter was employed to determine if high or low *eFGFs* expression and methylation in COAD had any predictive value for patient survival (Fig. 5A-E). Those patients who had high levels of *FGF21* and *FGF23* methylation (log-rank p = 0.03, 0.05 and hazard ratio (HR) = 0.59, 0.63, respectively) and low levels of *FGF23* expression (log-rank p = 0.02, HR = 0.57) had a considerably worse OS (Fig. 5C-E). While neither expression nor methylation levels of *FGF19* didn’t affect OS of COAD patients (*P-value* > 0.05) (Fig. 5A and B).

**FIGURE 5.**
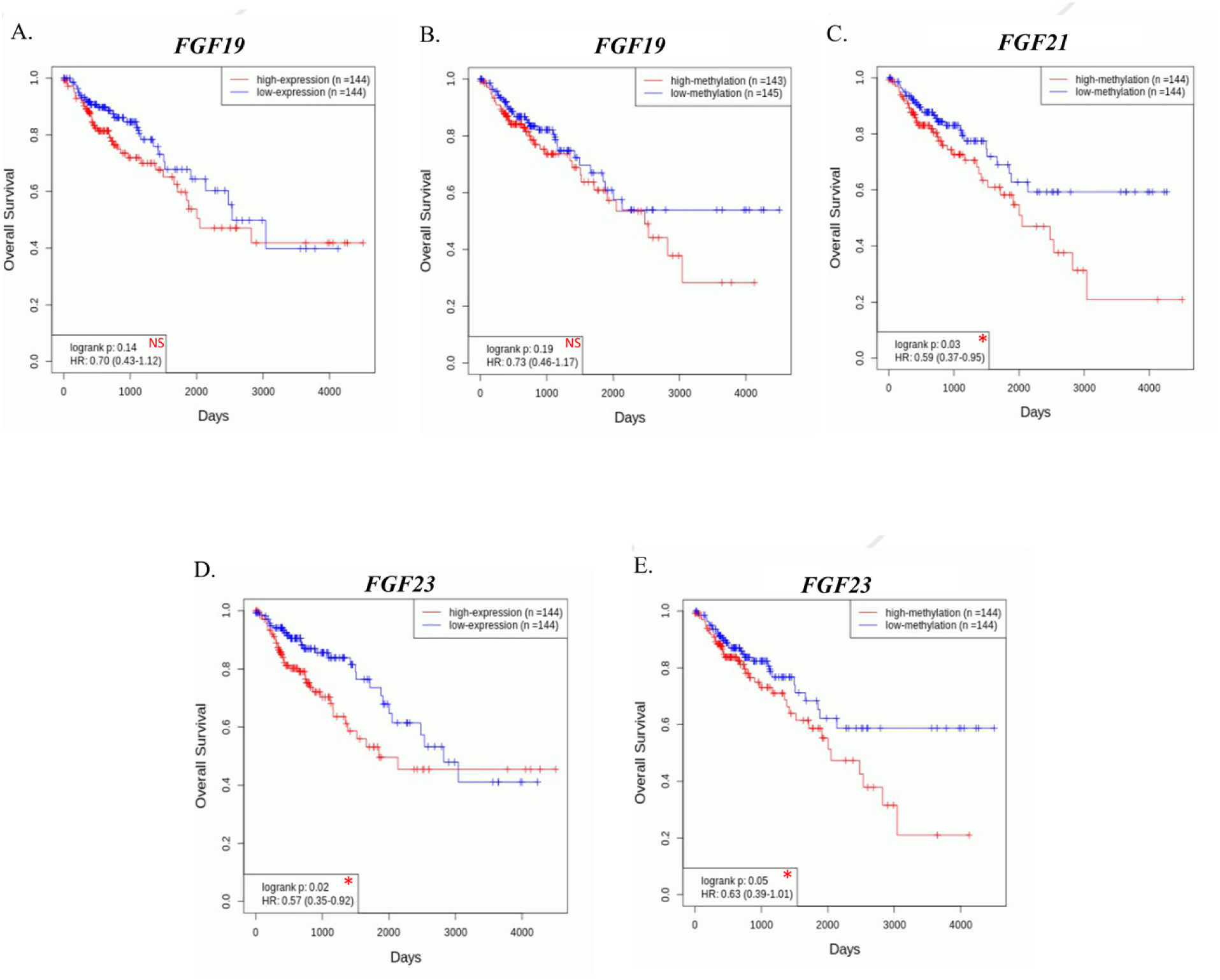
Prognostic value of endocrine *FGFs* (based on expression and methylation) in COAD patients via the ONCODB database. Overall survival curves for *FGF19* (A, B), *FGF21* (C), and *FGF23* (D, E) in COAD patients. (NS: *log-rank p* > 0.05, *: *log-rank p* < 0.05). NS; Non-significant. HR: Hazard ratio

### 3.5. Genetic Alteration in the eFGF Genes among CRC Patients

Using the cBioPortal database, we could measure the frequency of genetic alterations in the *eFGF* genes among COAD patients. Figure 6A demonstrates that eighteen COAD patients (8.18%) had varieties of significant alterations in the *eFGF* family members, such as mutation, amplification, transcriptional dysregulation, or multiple alterations. To be more precise, 4%, 4%, and 2.3%, respectively, of COAD patients had a genetic alteration in the *FGF19*, *FGF21*, and *FGF23* genes (Fig. 6B). With the goal of deciphering the mutational landscape of COAD across protein domains, we conducted an in-depth analysis, which revealed *F188L*, *R163H*, and *A72S* missense mutations for *FGF19*, *FGF21*, and *FGF23*; respectively (Fig. 6 C-E).

**FIGURE 6.**
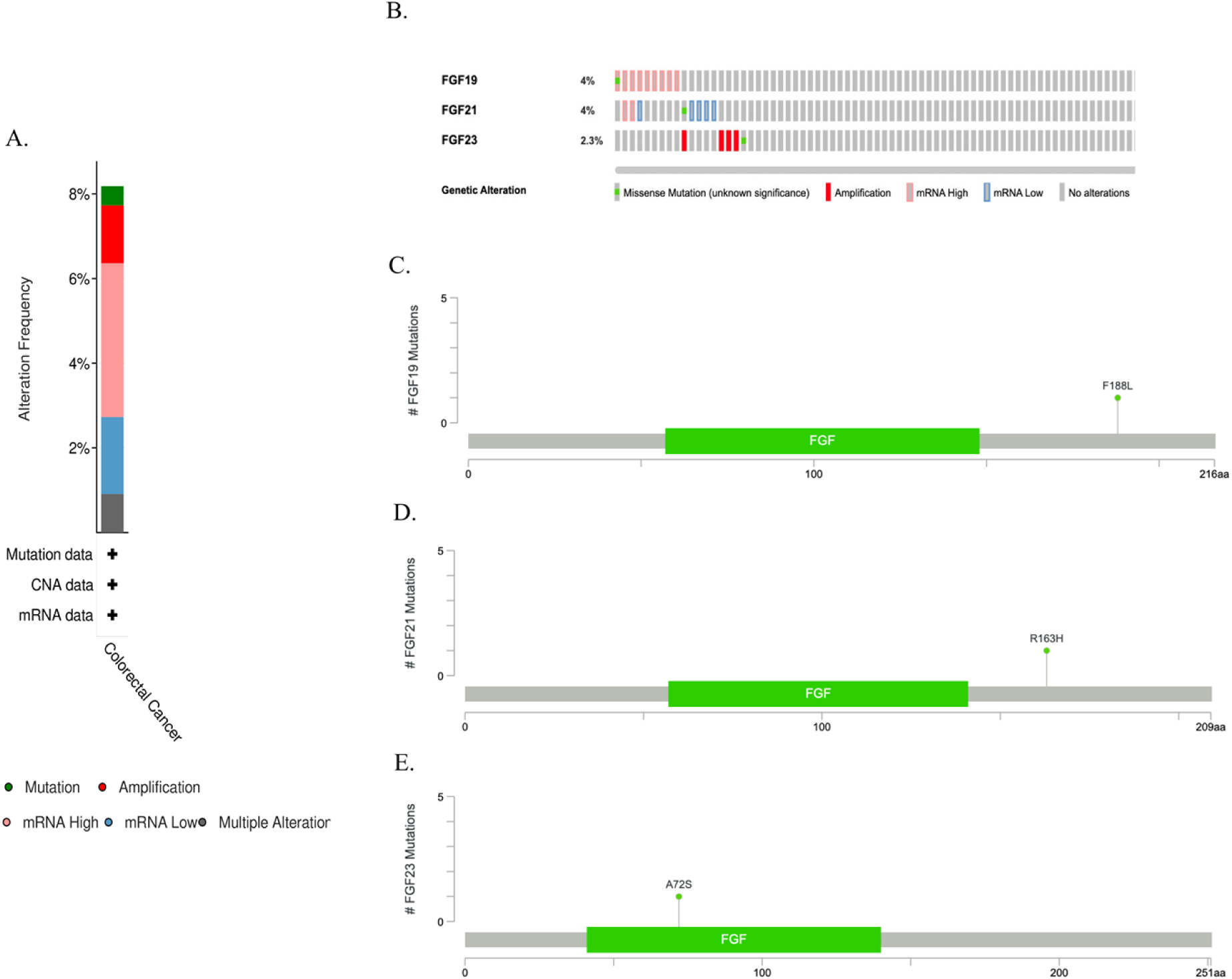
Alteration frequency of endocrine *FGFs* in COAD study based on cBioPortal database. **(A)** Summary of alterations in *eFGFs* in COAD. **(B)** OncoPrint visual summary of *eFGF* family members’ alterations. **(C-E)** Mutation diagram of *FGF19* **(C)**, *FGF21* **(D)**, and *FGF23* **(E)** across protein domains. Green bullets are missense mutations. Data are derived from the Mutations tab of the cBioPortal database.

### 3.6. Co-Expression Genes, Protein-Protein Interaction Network Construction, and Functional Annotation of eFGFs in COAD

We postulated that *eFGFs*’ function in COAD could be intimately tied to that of its adjacent genes. In light of this, *eFGFs* co-expression genes were analyzed in COAD patients using the LinkedOmics database. Genes that were positively or negatively correlated with *FGF19* (Fig. 7A), *FGF21* (Fig. 7B), and *FGF23* (Fig. 7C) were represented by dark red dots and dark green pots in volcano plots (false discovery rate, |FDR|<0.01). Figures 7D-I also demonstrated heat maps of the positive and negative correlations between *eFGF* family members and fifty significant gene sets. By the use of the STRING website, we constructed a PPI network to better assess the interactions for *eFGFs* (Fig. 7J-L). Finally, we performed KEGG enrichment analysis on *eFGF* genes. According to the findings, *eFGFs* have a function in the regulation of the actin cytoskeleton and, more notably, in Ras signaling, PI3k-Akt signaling, Rap1 signaling, and cancer pathways (Fig. 7M-O).

**FIGURE 7.**
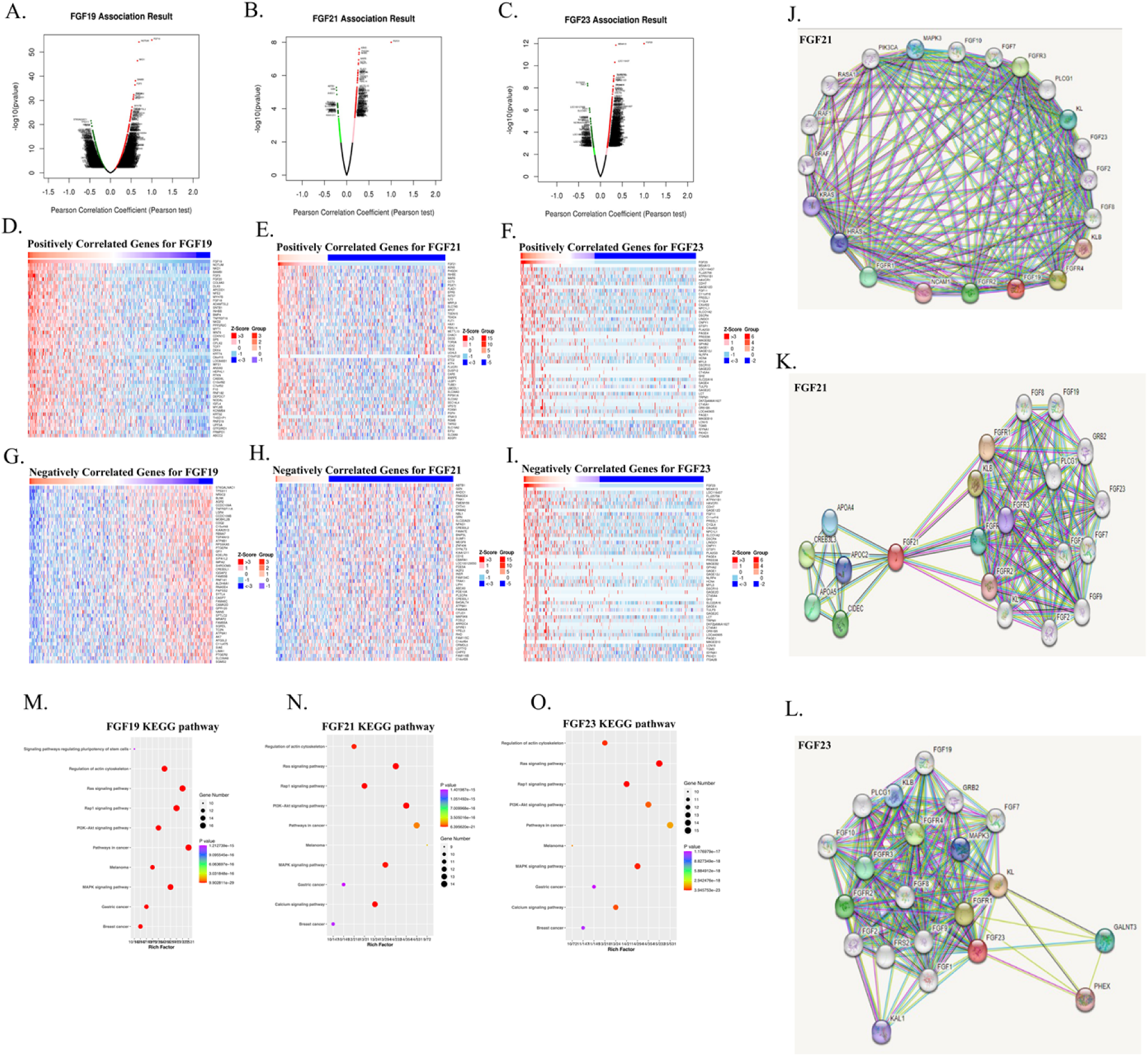
Co-expression genes of endocrine *FGFs* in COAD (LinkedOmics) and protein-protein interaction (PPI) network construction (STRING). (**A-C)** Volcano Plot demonstrating the positively and negatively correlated genes with *eFGFs*. **(D-I)** Heat maps show the top 50 significant genes positively correlated with *FGF19* (D), *FGF21*(E), and *FGF23* (F) and negatively correlated with *FGF19* (G), *FGF21* (H), and *FGF23* (I) in COAD. **(J-L)** Predicting the PPI network of *FGF19* (J), *FGF21* (K), and *FGF23* (L) using STRING. **(M-O)** KEGG pathway analysis of *FGF19* (M), *FGF21* (N), and *FGF23* (O).

### 3.7. The eFGF family members’ small molecule pathway Construction

We used SMPDB analysis to identify the human small molecule pathways in which the *eFGF* family members might be involved. We found that *FGF19* is a part of the bile acid indirect signaling pathway (Fig. 8). Enterocytes, which are epithelial cells of the small intestine, absorb bile acids, where they can activate the nuclear receptor farnesoid X receptor (FXR). As a result, *FGF19* is produced and released from the enterocyte and transported to the portal vein. Although most *FGF19* is transported to the liver, some *FGF19* instead enters systemic circulation, where it can penetrate the blood-brain barrier and interact with its receptors in the brain. A transmembrane protein called beta-klotho promotes the binding of *FGF19* with the receptor to form a stable complex. *FGFR* signaling probably contributes to the metabolism of glucose and energy. Nevertheless, there were no available data for *FGF21* and *FGF23* in the SMPDB database.

**FIGURE 8.**
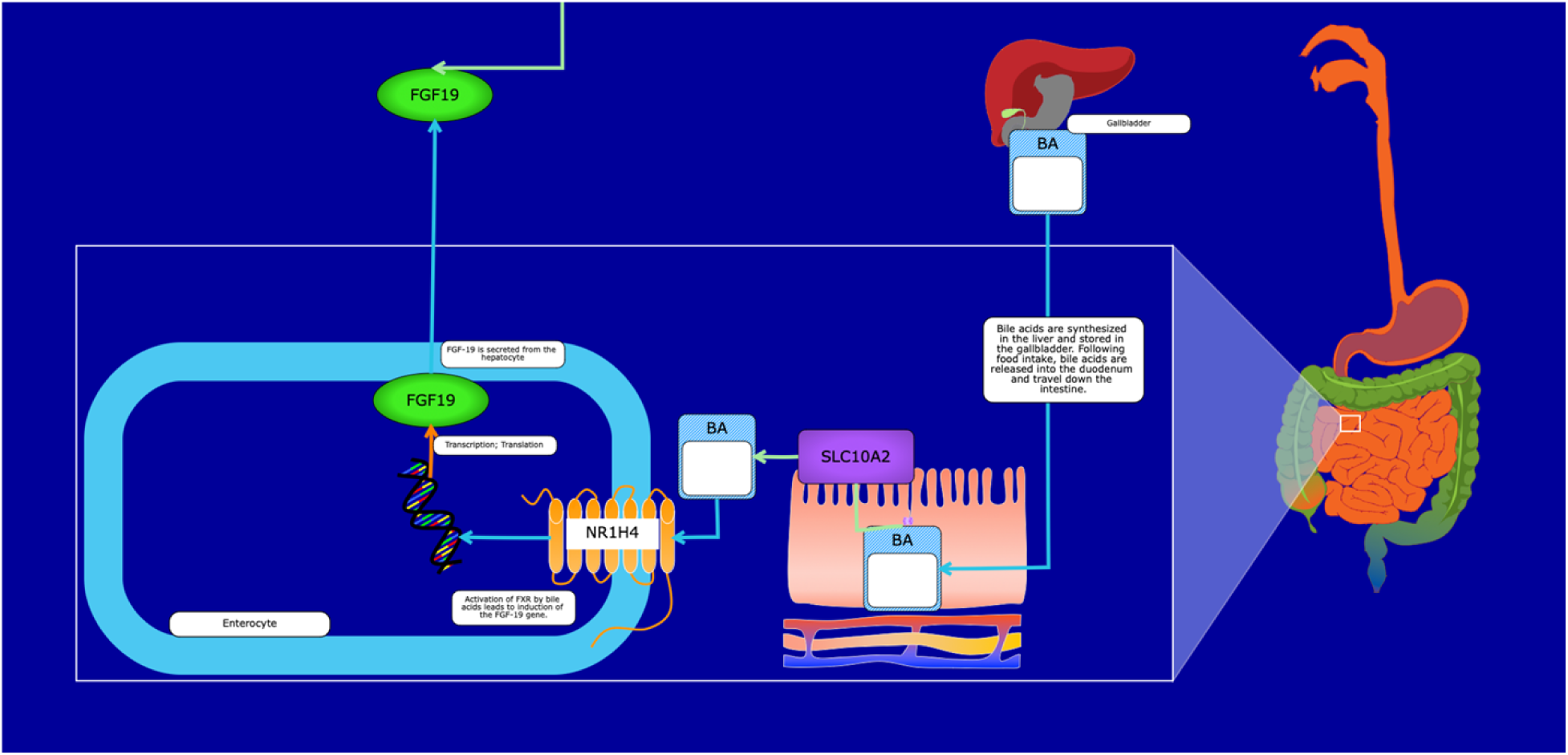
The use of the SMPDB database to analyze the small molecule pathways. The small molecular pathway of *FGF19* in humans.

### 3.8. Gene regulatory network

Transcriptional regulation is a fundamental biological process, and considerable effort has been expended worldwide to dissect it and identify its key characteristics to comprehend how its molecular components work together to regulate gene expression levels (38). Gene Regulatory Networks (GRNs) are potent instruments for describing and computationally reconstructing the intricate network of interactions underlying the transcriptional regulation of gene expression. GRNs are topological maps that represent and predict molecular entity relationships. On occasion, GRNs consist of regulatory interactions between a TF and its target genes, as well as between noncoding RNAs and target genes as a ceRNA network (39). The TF and ceRNA regulatory networks as GRNs were the primary focus of our investigation.

### 3.9. eFGF genes-Related Transcription Factors Regulatory network

TFs are vital components of signal transduction expressed in all human cells. They are generally regarded as the terminal effectors of signaling pathways in cells. As such, deregulating of a few TFs could profoundly impact overall gene expression profiles. Recent studies have underlined the importance of TFs in cancers (40–42), and revealed the crucial functions of TFs in tumor initiation, progression, and metastasis (43). To expound on the role of TFs in the regulation of CRC, we queried the potential TFs for *FGF19, FGF21,* and *FGF23* in colon tissue, and 12, 24, and 4 TFs were identified, respectively (Table S1). Figure 9A shows the TFs-Genes network and two TFs (CTCF and RAD21) regulate all members of *eFGF* family. Next, we investigated the expression of CTCF and RAD21 in CRC tissues and interestingly, these TFs were significantly overexpressed in tumor tissues compared to corresponding tissues (Fig. 9B).

**FIGURE 9.**
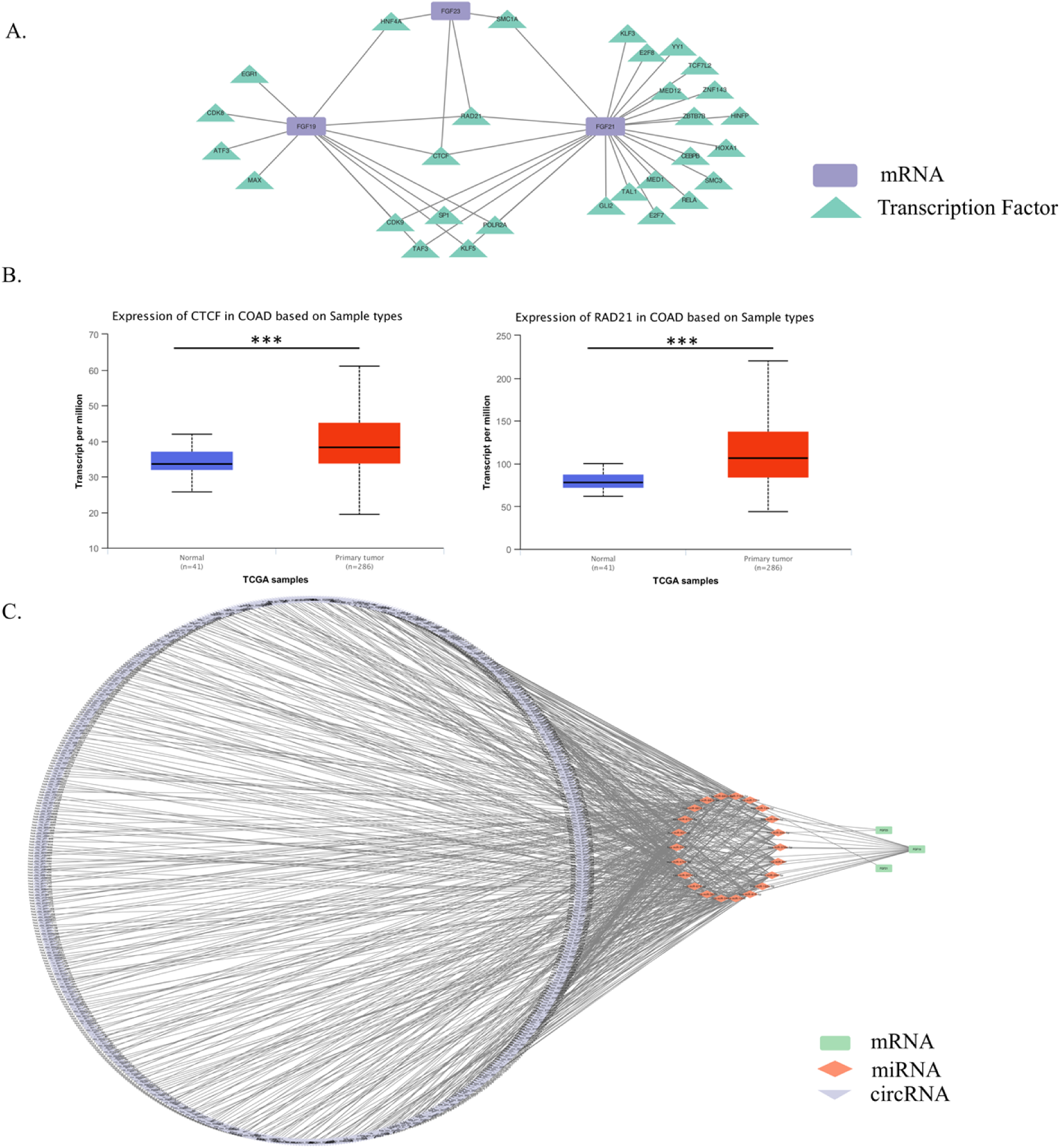
Endocrine *FGFs* Regulatory network prediction. **(A)** TF-Genes regulatory network prediction. **(B)** Expression of CTCF and RAD21 in COAD tissue compared to normal tissues. **(C)** The ceRNA regulatory network of *eFGF* family members. TF: Transcription factor; circRNAs: circular RNAs; miRNAs: micro-RNAs. (NS: *P*-value > 0.05, *: *P*-value < 0.05, **: *P*-value < 0.01). NS: Non-significant.

### 3.10. Prediction of eFGF family members’ mRNAs – miRNAs – circRNAs regulatory networks

The competing endogenous RNA is a novel regulatory mechanism in cancer, which is attracting widespread attention, and emerging evidence demonstrates that the ceRNA hypothesis has played a role in molecular biological mechanisms of occurrence and development of various cancer, including hepatocellular carcinoma, gastric, colorectal, and pancreatic cancer (44–47). To better understand the role of circRNAs and miRNAs in the ceRNA network of *eFGFs*, we established a circRNA–miRNA–mRNA (ceRNA) network. As the first step, we identified a total of nineteen, one, and two miRNAs targeting *FGF19, FGF21,* and *FGF23* mRNA, respectively (Table S2). Further, we retrieved circRNAs data that interact with miRNAs from the CircBank online database (Table S3). Finally, the ceRNA networks based on 660 circRNA nodes, twenty-two miRNA nodes, and three mRNA nodes were constructed and visualized (Fig. 9C and Table S4).

### 3.11. Drug - eFGF genes interaction

The DGIdb database was used to identify the drugs interacting with the e*FGF* family members. Burosumab was identified as an *FGF23* inhibitor with an interaction score of 247.32. Burosumab is a humanized monoclonal antibody against *FGF23* that has been authorized for the treatment of tumor-induced osteomalacia (48). However, drug interactions for *FGF19* and *FGF21* were not indicated by DGIdb.

### 3.12. Validation Analysis of eFGFs’ Pattern in CRC and Neoplasia by Real-time PCR

Then to corroborate the findings of the bioinformatics study, the expressions of *eFGFs* in 54 Polyps, 30 CRC, and 30 normal healthy tissues were identified by qRT-PCR. We found that the mRNA expression of *FGF21* was significantly higher in polyps than in CRC (*P-value* <0.008) or normal tissues (*P-value* <0.01) (Fig. 10A). Polyp tissues also had higher expression levels of *FGF19* and *FGF23* than tumor or normal tissues; however, these associations were not statistically significant (*P-value* > 0.05) (Fig. 10B and C). Moreover, the *FGF19*, *FGF21*, and *FGF23* expressions were shown to be elevated in CRC samples compared with normal tissue. But parallel to the prior results, except for *FGF21* (*P-value* = 0.045), the *FGF19* and *FGF23* associations were not statistically significant (*P-value* > 0.05) (Fig. 10A-C).

**FIGURE 10.**
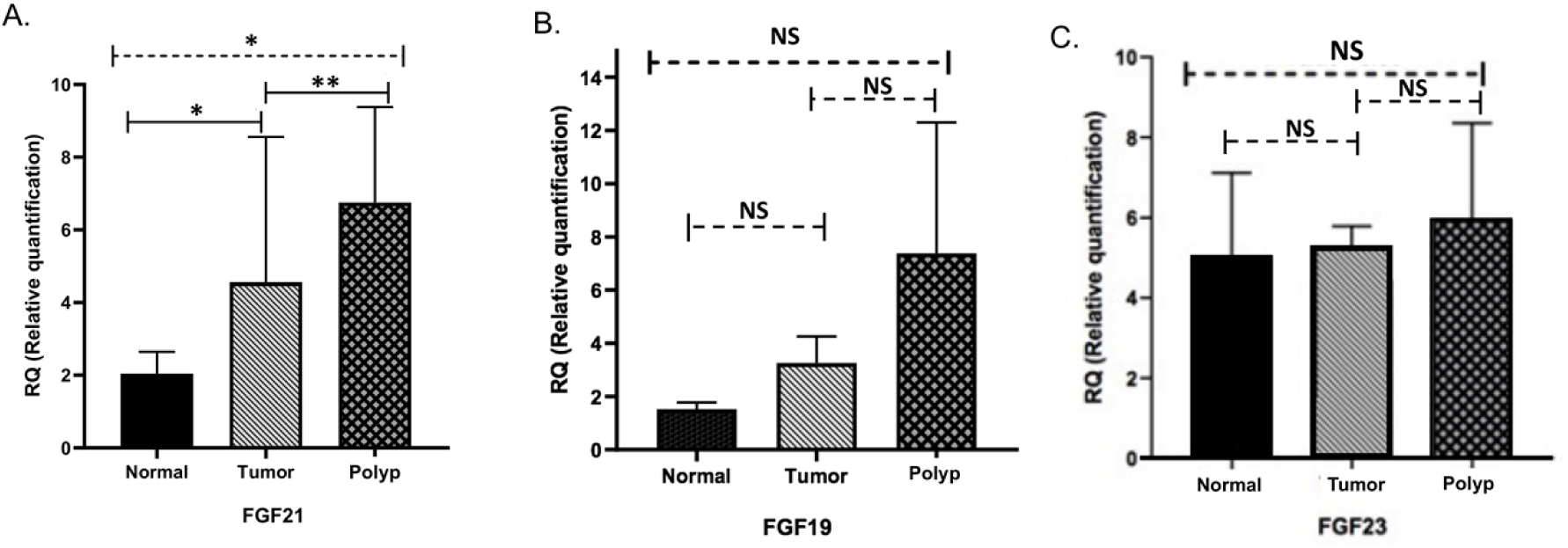
Relative mRNA expression of endocrine *FGF* between polyps and CRC and normal tissues. **(A)** Relative expression of *FGF21* between polyp and CRC (*P-value* < 0.008) and normal tissue (*P-value* < 0.01). **(B)** Relative expression of *FGF19* between polyp and CRC (*P-value* > 0.05) and normal tissues (*P-value* > 0.05). **(C)** Relative expression of *FGF23* between polyp and CRC (*P-value* > 0.05) and normal tissues (*P-value* > 0.05). (NS: *P*-value > 0.05, *: *P*-value < 0.05, **: *P*-value < 0.01). NS: Non-significant.

### 3.13. Association between eFGFs Expression and Overall Survival of CRC Patients’ Cohort

The log-rank analysis of our CRC patients’ cohort demonstrated that OS was significantly worse in patients with higher *FGF19* and *FGF21* expressions (*P-value* = 0.025 and 0.006; respectively) (Fig. 11A and B). Further multivariate analysis confirmed that *FGF19* and *FGF21* expression levels were independent prognostic indicators for OS of CRC patients (HR: 5.97 and 6.759; in order) (Table 1).

**FIGURE 11.**
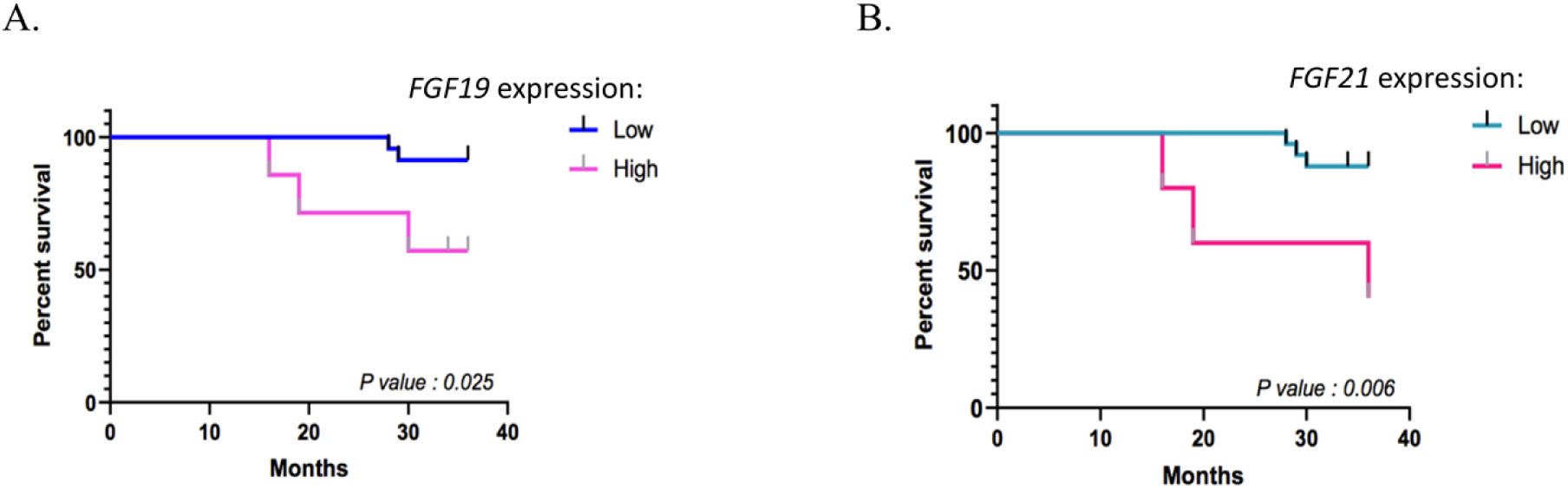
OS of patients with CRC based on *FGF19* and *FGF21* expression status. **(A, B)** CRC patients were equally divided into two groups based on *FGF19* (p-value:0.025) **(A)** and *FGF21* (p-value: 0.006) **(B)** mRNA levels, and then KM survival curves were employed for comparing OS between the two groups.

**Table 1.** Multivariate analysis for overall survival of CRC patients based on eFGF expression.

| Gene Name | HR | 95%CI of ratio | *P-value* |
| --- | --- | --- | --- |
| **FGF19** | 5.970 | 0.6695 to 53.24 | **0.025\*** |
| **FGF21** | 6.759 | 0.6205 to 73.62 | **0.0066\*** |
| **FGF23** | 2.454 | 0.4266 to 14.12 | 0.220 |

HR: Hazard Ratio, CI: Confidence Interval

### 3.14. Evaluation of eFGFs in Neoplastic Tissue as Potential Discriminating Factors Between High and Low-Risk Polyps

Given the importance of identifying high-risk polyps from low-risk ones in clinical practice, we assessed *eFGF* gene expressions in high-low-risk colon polyps. According to qRT-PCR findings in polyp tissues, the relative expression levels of *FGF21* and *FGF23* in AD were substantially higher than in HPP (*P-values < 0.02*) (Fig. 12 A and B, respectively). In light of these findings, it is plausible that these two genes contribute as potential distinguishing markers of high-risk polyps (3) from low-risk polyps (HPP).

**FIGURE 12.**
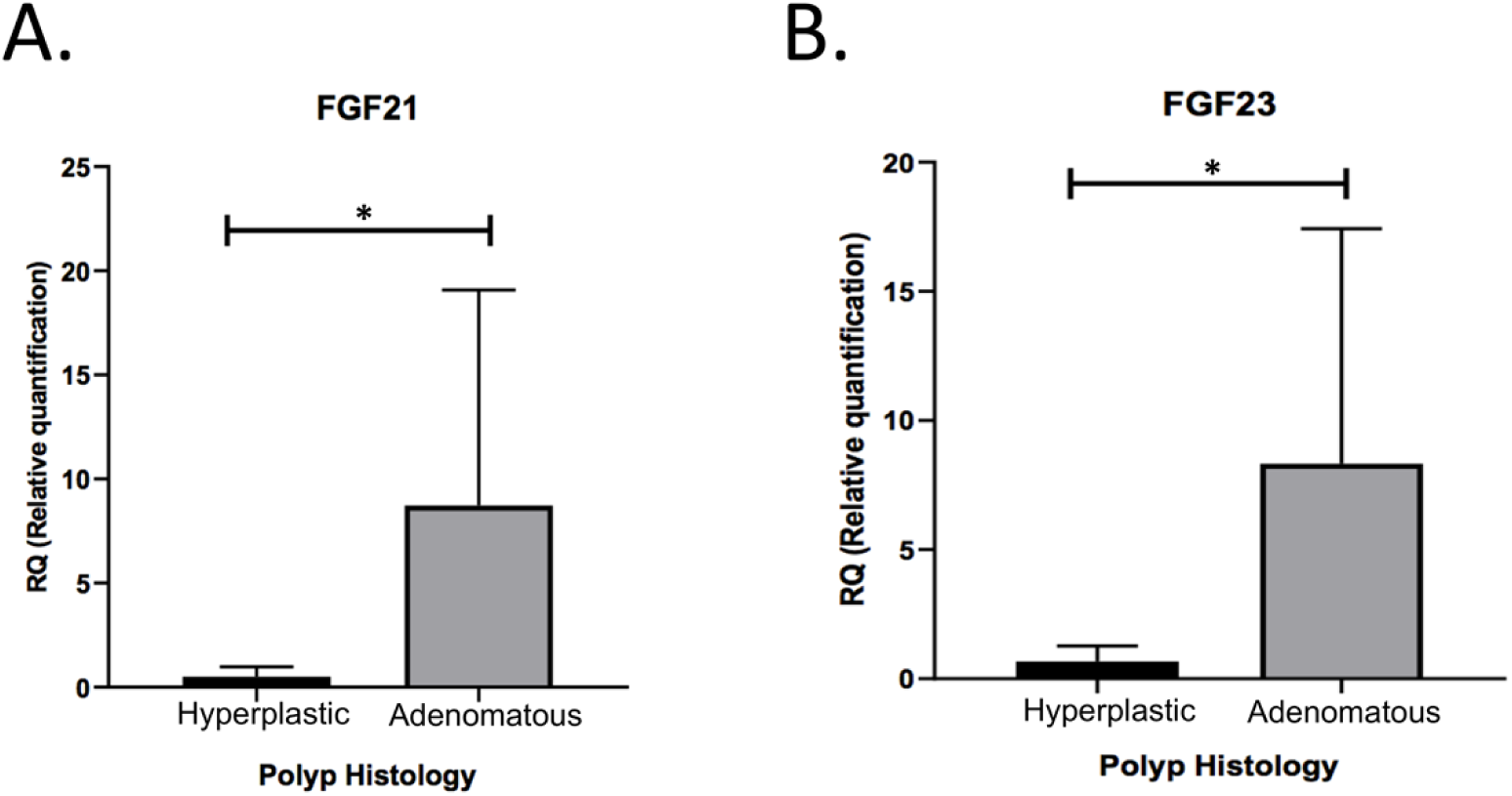
qRT-PCR analysis of *FGF21* and *FGF23* expressions in 26 adenomatous polyps and 28 hyperplastic polyps. **(A and B)** Expression levels of *FGF21* (*P-value* = 0.02) **(A)** and *FGF23* (*P-value* = 0.02) **(B)** were significantly elevated in adenomatous tissue compared to hyperplastic polyps. (* indicates *P*-value < 0.05)

### 3.15. Association between Expression Levels of eFGSs and Clinicopathological Features in Cohorts of Polyp and CRC Patients

Eventually, we assessed the correlation between *eFGFs* expression and clinical characteristics using expression levels obtained from qRT-PCR data of two independent cohorts of colon polyp and CRC patients (Tables 2 and 3). In our colon polyp cohort, we found that *FGF23* expression level was significantly associated with the gender of patients (*P-value* = 0.03), polyp size (*P-value* = 0.03), and anatomical site of polyps (*P-value* = 0.04) (Table 2). Likewise, among the CRC patients’ cohort, expression of *FGF19* was significantly related to tumor size (*P-value* = 0.02) and TNM stage (*P-value* = 0.03). *FGF21* was associated with the TNM stage (*P-value* = 0.012) and pathologic differentiation (*P-value* = 0.04) (Table 3).

**Table 2.** Correlation between endocrine FGFs expression and clinicopathological characteristics of patients with the polyp.

| Factor | FGF19 |  | FGF21 |  | FGF23 |  |
| --- | --- | --- | --- | --- | --- | --- |
|  | (SD) ± Mean of RQ | *P-value* | (SD) ± Mean of RQ | *P-value* | (SD) ± Mean of RQ | *P-value* |
| **Gender**<br>Female<br>Male | 1.84±1.67<br>4.10±2.91 | 0.94 | 4.56±3.09<br>12.05±9.49 | 0.75 | 1.37±1.70<br>10.07±11.67 | **0.03\*** |
| **Age**<br>>50<br>≤50 | 3.99±2.65<br>1.98±1.88 | 0.89 | 11.53±8.68<br>5.08±3.26 | 0.60 | 10.30±8.63<br>2.90±1.17 | 0.29 |
| **Polyp size**<br>>5 cm<br>≤5 cm | 7.41±4.54<br>0.86±0.93 | 0.80 | 12.36±10.83<br>4.56±3.09 | 0.44 | 10.07±11.67<br>1.20±1.64 | **0.03\*** |
| **Histology**<br>Hyperplastic<br>Adenomatous | 0.49±1.08<br>5.12±4.00 | 0.76 | 0.47±0.50<br>10.34±8.72 | **0.02\*** | 0.60±0.67<br>9.10±8.32 | **0.02\*** |
| **Differentiation**<br>High grade<br>Low grade | 4.57±3.61<br>5.04±2.70 | 0.85 | 2.13±1.31<br>11.54±9.32 | **0.02\*** | 11.07±8.43<br>6.78±5.37 | 0.94 |
| **Location**<br>Left<br>Right | 1.84±1.67<br>6.95±4.16 | 0.94 | 4.56±3.09<br>12.05±9.49 | 0.75 | 1.28±1.97<br>9.60±10.85 | **0.04\*** |

SD: *standard deviation*, RQ: Relative quantification

**Table 3.** Correlation between endocrine FGFs expression and clinicopathological features of CRC patients.

| Factor | FGF19 |  | FGF21 |  | FGF23 |  |
| --- | --- | --- | --- | --- | --- | --- |
|  | (SD) ± Mean of RQ | *P-value* | (SD) ± Mean of RQ | *P-value* | (SD) ± Mean of RQ | *P-value* |
| **Gender**<br>Female<br>Male | 4.30 ±2.90<br>6.39 ±3.52 | 0.47 | 4.47 ±1.96<br>0.68 ±0.66 | 0.97 | 6.91 ±4.14<br>8.37 ±4.87 | 0.32 |
| **Age**<br>>50<br>≤50 | 1.66±1.80<br>9.19±6.64 | 0.88 | 0.97 ±0.68<br>5.25 ±2.49 | 0.35 | 7.95± 4.79<br>7.61 ±4.50 | 0.56 |
| **Tumor size**<br>>5 cm<br>≤5 cm | 7.99±6.81<br>1.08±1.24 | **0.02\*** | 0.80±0.57<br>1.60±1.10 | 0.19 | 10.65±8.16<br>2.42±1.88 | 0.09 |
| **TNM stage**<br>I and II<br>III and IV | 5.11 ±2.55<br>6.20 ±4.76 | **0.03\*** | 3.79 ±1.92<br>0.25 ±0.33 | **0.012\*** | 8.89 ±5.61<br>2.57 ±2.19 | 0.72 |
| **Differentiation**<br>Poor<br>Well and Mod | 1.64 ±1.93<br>6.32 ±3.73 | 0.68 | 0.41 ±0.34<br>3.53 ±1.59 | **0.04\*** | 9.59 ±5.14<br>6.80 ±4.31 | 0.49 |
| **Vascular invasion**<br>Yes<br>No | 4.44 ±3.19<br>6.44 ±3.31 | 0.53 | 1.23 ±0.89<br>3.98 ±1.51 | 0.89 | 9.59 ±5.93<br>2.72 ±2.16 | 0.39 |
| **Lymph node invasion**<br>Yes<br>No | 4.63 ±2.96<br>7.75 ±4.07 | 0.46 | 3.47 ±1.48<br>0.39 ±0.52 | 0.89 | 7.69 ±4.32<br>7.60 ±4.59 | 0.75 |
| **Anatomical site**<br>Left<br>Right | 6.87 ±4.08<br>2.04 ±2.01 | 0.89 | 3.75 ±1.53<br>1.25 ±0.76 | 0.27 | 7.88 ±4.60<br>7.33 ±4.08 | 0.84 |

SD: standard deviation

## 4. Discussion

In our study, we first examined the expression of endocrine *FGF* family members in COAD tissues compared with their NAT and healthy individual colon samples. Second, we provide evidence that *eFGFs* expression and methylation levels are associated with different clinicopathological characteristics of COAD and also determined that expression or methylation of some of these genes are related to the OS of COAD patients. What’s more, we identified co-expressed genes and scrutinized functional analysis and molecular pathways, which were discussed in more detail later in this assay. Finally, we verified the expression of *eFGFs* in CRC tissue, colon polyps, and normal tissue samples by GSE and qRT-PCR. Our experimental study showed that the expression of *FGF21* in colon polyps was considerably higher than that in CRC or normal tissues.

Endocrine *FGF* family members contribute to inter-organ endocrine signaling axes that are essential for whole-body homeostasis, since they regulate bile acid, glucose, and lipid metabolism (49). In essence, *eFGFs* have two distinct features, a growth factor and a metabolic regulator. Hence, looking into *eFGFs* may illuminate the molecular connection between cancer development and metabolism (50). Accordingly, significant variations in *eFGF* expression have been linked to the development of various diseases, specifically cancers (51–53).

Based on our findings, *eFGFs* transcriptional expressions were identified as being higher in COAD tissues than in NAT tissues. We found that a series of investigations supported the oncogenesis hypothesis of *eFGFs*. Tumorigenesis and the development of CRC and urothelial carcinoma in vitro and in vivo were both linked to *FGF19* overexpression (53, 54).

Proteomic profiling by Harlid et al. (55) in the pre-diagnostic plasma of 138 CRC patients showed an association between high *FGF21* levels and an increased risk of CRC. *FGF23* has been linked to osteogenic osteomalacia in colon and urothelial cancer (22, 53). Thus, these data suggest that *eFGFs* may serve as potential biomarkers for several cancers.

Subsequently, both bioinformatics and experimental data determined a significant correlation between *FGF19* expression levels and COAD tumor stages. However, this link did not extend to other *eFGFs* (*FGF21* and *FGF23*). In a similar vein, Motylewskaa et al.(56) showed that *FGF19* overexpression in gastric cancer is linked to the TNM stage, depth of invasion, and lymph node metastasis. Using qRT-PCR, we further demonstrated that increased levels of *FGF21* were related to both the histopathological differentiation and the TNM stage of the CRC patients’ cohort. In line with our findings, Cymbaluk-Poska et al. (17) demonstrated a correlation between the serum levels of *FGF21* and the endometrial Cancer clinical stage. Circulating *FGF21* levels were also associated with an elevated risk of CRC in all stages, according to Qian et al. (51). In our analysis, although the *FGF23* mRNA expression was significantly elevated in more advanced stages compared to normal colon tissue, there was no correlation between the expression level of *FGF23* and the various stages of COAD cancer, according to the GEPIA2 database.

Using bioinformatics tools, we also conducted the subgroup analysis for the promotor methylation status of *eFGF* genes. The primary tumor tissues, tumor stages, and tumor histology were demonstrated to correlate significantly with the *FGF19* and *FGF21* promotor hypermethylation when compared to normal tissues. In exactly the opposite manner, the *FGF23* promotor was hypomethylated in all mentioned characteristics.

In our predictive analysis based on expression and methylation datasets from OncoDB, we were able to reveal that COAD patients with hypermethylated *FGF21*, *FGF23*, or lower mRNA expression of *FGF23* had shorter OS. According to qRT-PCR results in our independent CRC cohorts, patients with higher expressions of *FGF19* and *FGF21* had a worse prognosis than those with lower expressions. Overall, these findings indicated that *FGF* family members, particularly *FGF21* expression and methylation levels, are potential and promising prognostic indicators for COAD. A recent study has shown that endometrial cancer patients with high expression for FGF21 have a worse prognosis (17).

Premalignant cells often require more nutrients and energy to maintain fast proliferation because of their high metabolic rate (57, 58). Recent research has demonstrated that metabolic reprogramming is governed by the expression of oncogenic pathways such as Ras, PI3k-Akt, P53, and WNT signaling pathways, which are also strongly linked with the onset and development of CRC (59, 60). These data are consistent with our pathway enrichment analysis results. Based on the KEGG pathway, we identified that *eFGF* family members are mainly involved in Ras, PI3k-Akt, Rap1, and cancer pathways, besides regulation of the actin cytoskeleton.

With the development of cancer, nutrient deprivation may rapidly trigger *eFGFs* secretion (51). Nicholes et al. (61) have shown that *FGF19* may act as a promotor of cancer in the hepatocellular carcinoma (HCC) transgenic model. Given the close link between dysplastic cells and metabolism, it is rational to infer that *eFGF* might also be related to the initiation of colorectal carcinogenesis. Herein, for the first time, we provide preliminary evidence of the importance of *eFGFs* as early CRC markers. The gene expression analysis revealed higher *FGF19, FGF21,* and *FGF23* in the colon polyps than in normal tissues (*p-value =* ns, <0.013, ns; respectively). Compatible with these findings, higher circulating levels of *FGF21* increased the risk of metachronous colorectal adenomas, especially in the older population (18)

Colonoscopy is the gold standard for detecting colon cancer at an early stage and for following-up patients after a polypectomy (62, 63). Even with a colonoscopy, though, it is not always possible to distinguish high-risk from low-risk polyps based on their appearance (64, 65). According to a meta-analysis that included 43 studies involving more than 15,000 tandem colonoscopies, the adenoma miss rate was reported to be 26% (66). Our qRT-PCR data indicated that AD, as opposed to HPP, had significantly higher expression levels of *FGF21* and *FGF23*. It has also been claimed that colonoscopy is less effective in preventing proximal than distal colon cancer (67). Perhaps due to distinct oncogenesis mechanisms, right-sided colon cancers are more likely to be detected at an advanced stage and have an unfavorable prognosis than left-sided tumors (68). The current investigation found that right-sided polyps expressed considerably more significant levels of *FGF23* than left-sided polyps. As a result, these two genes have the potential to serve as biomarkers for discriminating high-risk from low-risk polyps. To our knowledge, our study provides the first demonstration of the clinical significance of *eFGF* in colorectal neoplasia, though the results need to be further replicated.

Some strengths and limitations of our study should be taken into account. The principal strength of the present investigation is a population-based study that includes CRC and Colon Polyps cohorts from screening settings, along with a comprehensive and systematic bioinformatics analysis of *eFGFs* in COAD. Even though the study populations were large in total, the number of participants with polyps, particularly those with AD, remained fairly limited. Nevertheless, replication in larger studies, ideally with repeated longitudinal measurements of *eFGF* family members, is needed to more fully understand the potential role of these genes in colorectal carcinogenesis. Besides, we have only experimentally validated the expression of *eFGFs* at the RNA level; further confirmation of *eFGFs* at the protein or methylation levels is highly recommended.

Consequently, our research sheds light on the potential biological mechanism of endocrine *FGFs*, which may lead to providing promising biomarkers for the diagnosis and prognosis of CRC patients as well as a novel treatment strategy. These preliminary results also open the door for the speculation that *FGF* family members are implicated in the first steps of CRC carcinogenesis.

## 5. Abbreviation

AD: Adenomatous polyp
AUC: Area Under ROC Curve
cDNA: complementary DNA
ceRNA: competing endogenous RNA
COAD: Colon adenocarcinoma
CRC: Colorectal cancer
DEB-TACE: Drug-eluting bead trans-arterial chemoembolization
eFGF: endocrine Fibroblast growth factors
FGFs: Fibroblast growth factors
FXR: Farnesoid X receptor
GRN: Gene Regulatory Network
HCC: Hepatocellular carcinoma
HPP: Hyperplastic polyp
HR: Hazard ratio
NAT: Normal Adjacent Tissues
OS: Overall survival
PPI: Protein-Protein Interaction
qRT-PCR: Real-Time Polymerase Chain Reaction
RNA-Seq: RNA sequencing
ROC curve: Receiver Operating Characteristic curve
TF: Transcription factor
TNM: Tumor-node-metastasis

## 6. Author Contributions

Conceptualization, Methodology, and Project Administration: ENM and ZS; In Silico Data Collection and Analysis: MP, PJ, and ZS; Experimental Design: ENM, MH, LR, EDA, and MJ. Experimental Data Screening and Data Collection: LR, MSN, BK, and MP; Draft Writing: MP and LR; Draft Review and Editing: ENM, ZS, LR, and MP; Thesis Guidance and Review: HAA, ENM, ZS, and MH. All Authors Approved the Final Version to be Published; They All Agreed to be Accountable for All Aspects of the Work.

## Acknowledgments

We thank everyone who provided support for this study.

## 7. Conflicts of Interest

The authors declare that they have no conflicts of interest.

## 8. Funding

We thank Taleghani Hospital, Shahid Beheshti University of medical sciences, for the financial support (code: 987). This article has not received sponsorship for publication.

## 9. Ethical Approval Statement

All subjects participated voluntarily and received a small compensation. The participants provided their written informed consent to participate in this study. the research was approved by the Ethics Committee of Taleghani Hospital at Shahid Beheshti University of Medical Sciences (protocol number: IR.SBMU.RIGLD.REC.1396.180).

## 10. Data Availability Statement

The experimental data supporting this study’s findings are not openly available due to reasons of sensitivity and are available from the corresponding author upon reasonable request. All gene expression data are available in the TCGA database, and the raw data of signaling pathways are available in KEGG. The datasets presented in this study can be found in online repositories. The names of the repository/repositories and accession number can be found in the article.

## Notes

### Competing Interest Statement

The authors have declared no competing interest.

